# Nanopore sequencing of full-length *BRCA1* mRNA transcripts reveals co-occurrence of known exon skipping events

**DOI:** 10.1101/169755

**Authors:** Lucy C de Jong, Simone Cree, Vanessa Lattimore, George AR Wiggins, Amanda B. Spurdle, kConFab Investigators, Allison Miller, Martin A Kennedy, Logan C Walker

## Abstract

Laboratory assays evaluating the effect of DNA sequence variants on *BRCA1* mRNA splicing may contribute to classification by providing molecular evidence. However, our knowledge of normal and aberrant *BRCA1* splicing events to date has been limited to data derived from assays targeting partial transcript sequences. For the first time, we resolve the exon structure of whole *BRCA1* transcripts using MinION nanopore sequencing of long-range PCR amplicons. Our study identified 32 *BRCA1* isoforms, including 18 novel isoforms which comprised skipping of multiple contiguous and/or non-contiguous exons. Furthermore, we show that known *BRCA1* exon skipping events, such as Δ(9,10) and Δ21, can co-occur in a single transcript, with some isoforms containing four or more alternative splice junctions. Our results highlight complexity in *BRCA1* transcript structure that has not previously been described. This finding has key implications for predicting translation frame of splicing transcripts, important for interpreting the clinical significance of spliceogenic variants. Future research is warranted to quantitatively assess full length *BRCA1* transcript levels, and to assess the application of nanopore sequencing for routine evaluation of potential spliceogenic variants.

## Introduction

Routine diagnostic screening for deleterious variants in the breast cancer susceptibility gene *BRCA1* is typically performed for individuals from suspected high-risk breast (and ovarian) cancer families to identify the genetic cause for their disease. However, an important practical issue associated with genetic testing is the identification of rare sequence variants with unknown clinical significance. Interpreting the clinical meaning of unclassified variants is a key challenge facing the future of genomic-based health initiatives.^1^

Multifactorial likelihood analysis is the most accepted approach for assessing cancer risk associated with unclassified *BRCA1* variants and has been successful in classifying hundreds of variants since it was developed^2, 3^ (http://brcaexchange.org/). However, the multifactorial likelihood model is limited by the amount of information available from the variant carrier (tumour histopathology), the family of the variant carrier (co-segregation, family history information) and additional information, such as co-occurrence with a pathogenic variant. Numerous studies have shown that the effect of a variant of unknown clinical significance on *BRCA1* mRNA splicing may contribute to classification by offering molecular evidence.^4-7^ Moreover, to classify variants using a combination of bioinformatic and *in vitro* splicing data, Spurdle et al. (2008) ^8^proposed 5-tier splicing classification (Class 5-pathogenic, Class 4-likely pathogenic, Class 3-uncertain, Class 2-likely not pathogenic, Class 1-not pathogenic) guidelines. These guidelines were subsequently improved after a multicentre study carried out by the international ENIGMA consortium.^6^

Determining which mRNA splice isoforms are abnormal and potentially deleterious can be challenging. The ENIGMA Splicing Working Group recently undertook a comprehensive analysis to characterize numerous ‘naturally occurring’ mRNA splice isoforms for *BRCA1* to aid in the interpretation of *in vitro* splicing assays.^9^ This study identified more than 60 *BRCA1* mRNA isoform events occurring in breast and/or blood cells. However, it remains unclear whether these individual splicing events can co-occur in the same *BRCA1* transcript, as PCR- and sequencing-based technologies used to assess splicing events typically interrogate only a fraction of the whole transcript(s). Pathogenic (or Class 5) variants that cause mRNA splicing changes are expected to disrupt protein function either through truncation or in-frame deletion of important regions of the encoded proteins. Using technologies that only examine a section of mRNA transcripts for variant classification may therefore lead to a misinterpretation of in-frame or out-of-frame splicing events.

DNA sequencing technology based on nanopore sequencing generates read lengths that greatly exceed those of more commonly used Sanger sequencing and massively parallel sequencing platforms. Moreover, nanopore sequencing has been demonstrated to characterise the complex exon structure of mRNA transcripts from genes expressing a large number of isoforms.^10^ To our knowledge, single-molecule sequencing technologies (MinION^11^ and PacBio^12^) that enable long sequence reads have yet to be employed to resolve the exon structure of whole *BRCA1* mRNA transcripts. In this study, we explored the utility of long-range reverse transcriptase (RT)-PCR with nanopore sequencing to identify novel *BRCA1* isoforms and the co-occurrence of known exon skipping events.

## Results and Discussion

### *PCR amplification of full*-*length* BRCA1 *cDNA*

Lymphoblastoid cell lines (LCLs) have been widely used as a cell model for evaluating *BRCA1* splicing changes.^9^ For this study, we assessed RNA from a healthy control LCL that was previously used for an international workshop, led by the ENIGMA Consortium, that compared mRNA splicing assay protocols between laboratories.^7^ To obtain full length *BRCA1* transcripts, we carried out long-range RT-PCR for *BRCA1* transcripts using primers targeting the 3’ end of exon 1 and the 5’ end of exon 24 to generate a 5.8kb amplicon (Supplementary Figure 1). Amplified products from repeat assays for a single LCL were visualised by gel electrophoresis, revealing a difference in patterns of amplicon sizes and suggesting variability in isoform selection and amplification during the PCR cycles (Figure 1; Supplementary Figure S2). PCR products that were consistent by size with a full-length *BRCA1* isoform (NM_007294.3: encoding the full-length BRCA1 protein) were observed in 28/47 PCR assays. To generate a sequencing catalogue of whole *BRCA1* transcripts, amplified products were pooled from all 47 PCR assays using cDNA synthesised from a single LCL RNA sample. A preliminary assessment of fragments from 10 PCR assays by Sanger sequencing confirmed *BRCA1* identity (Figure 1) prior to sequencing using the MinION.

**Figure 1.**
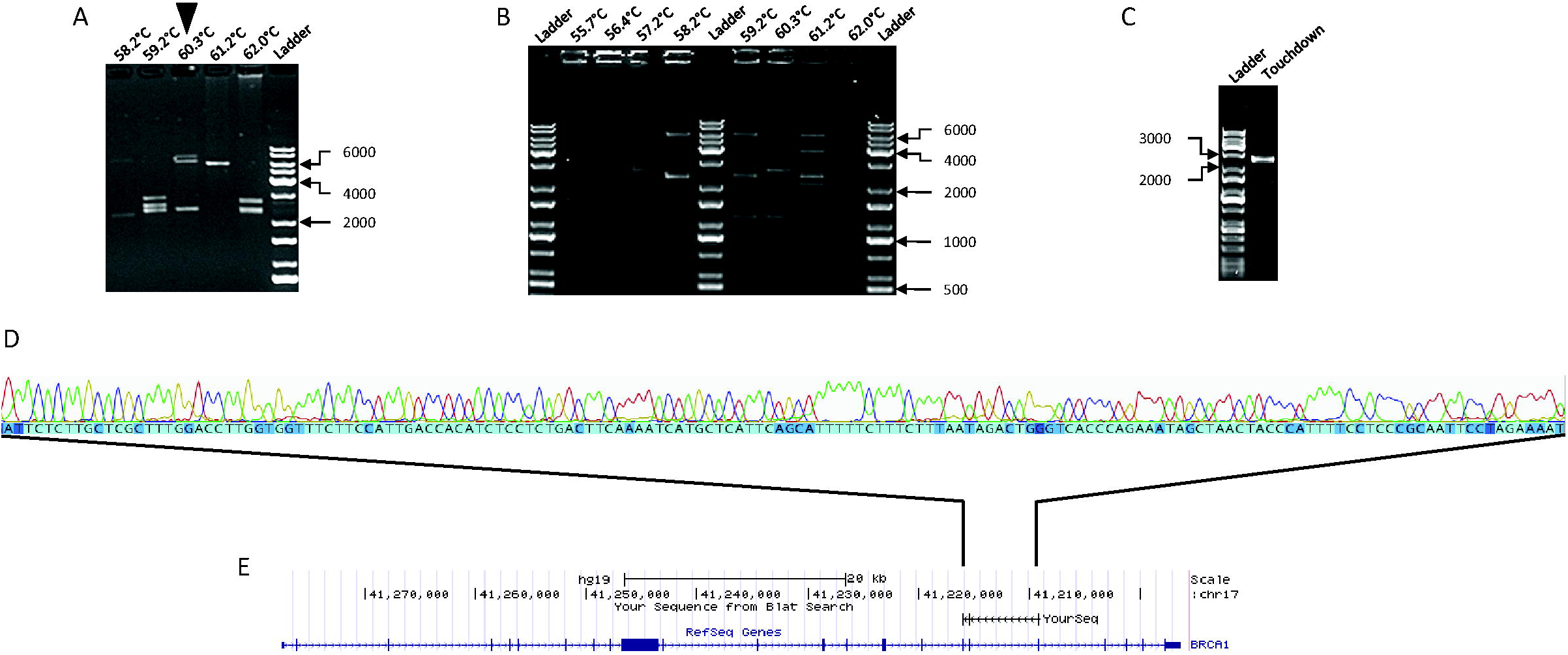
Example of fragments obtained from different long-range PCR assays from a lymphoblastoid cell line. A human lymphoblastoid cell line (LCL) derived from a female healthy control was cultured with cycloheximide to prevent nonsense mediated RNA decay (NMD), as previously described._5_ RNA was extracted from the cells using the RNeasy Mini Kit (Qiagen), according to the manufacturer’s instructions. cDNA synthesis was carried out using oligo(dT) primers (ThermoFisher Scientific Inc.) and Superscript^®^ III Reverse Transcriptase (ThermoFisher Scientific Inc.) according to the manufacturer’s instructions. Twenty μL of resulting cDNA mix was diluted 5-fold in H_2_0, and 3 μL of the final solution was used for each long-range PCR assay. Three different protocols were used to generate a pool of PCR amplicons for nanopore sequencing. PCR products were resolved in a 1% agarose gel using electrophoreses. **A**. Protocol 1: Reactions contained 1.3M betaine (Sigma-Aldrich), lx KAPA long-range buffer (KAPA Biosystems), 1.75mM MgCl_2_, 0.5μM of each primer (BRCA1_1F 5’-GCGCGGGAATTACAGATAAA-3’ and BRCA1_24pR 5’ -AAGCTCATTCTTGGGGTCCT-3’), 300μM of KAPA, 200μM dNTP mix, and 0.5 units of KAPA Long Range HotStart. Thermal cycling conditions were 94°C for 4 minutes, followed by 35 cycles of 94°C for 30 seconds, primer annealing at one of a range of temperatures (56.4oC to 62.5oC; Supplementary Figure 2) for 30 seconds, and 68°C for 12 minutes, with a final extension of 72°C for 12 minutes. Primer annealing temperature is indicated above each lane. Reference markers labelled for size in base pairs. Reference markers labelled for size in base pairs. **B**: Protocol 2: PCR reactions contained 1M betaine, 1x KAPA long-range buffer, 2mM of MgCl_2_, 0.7μM of each primer (BRCA1_1F and BRCA1_24pR), 200μM dNTP mix, and 0.5 units of KAPA Long Range HotStart. Thermal cycling conditions were 94°C for 2 minutes, followed by 35 cycles of 94°C for 30 seconds, 55.7oC to 62.5oC (Supplementary Figure 2) for 30 seconds, and 68°C for 7 minutes, before a final extension of 72°C for 7 minutes. Primer annealing temperature is indicated above each lane. Reference markers labelled for size in base pairs. **C**. Protocol 3. Reactions contained 1M betaine, lx KAPA long-range buffer, 2mM MgCl_2_, 0.7μM of each primer (BRCA1_1F and BRCA1_24pR), 200μM dNTP mix, and 0.5 units of KAPA Long Range HotStart. Thermal cycling conditions were 94°C for 2 minutes, then 8 cycles of 94°C for 30 seconds, 66°C for 30 seconds (decreasing 1°C each cycle), and 68°C for 7 minutes, followed by 30 additional cycles of 94°C for 30 seconds, 59oC for 30 seconds, and 68°C for 7 minutes, before a final extension of 72°C for 7 minutes. Primer annealing temperature is indicated above each lane. Reference markers labelled for size in base pairs. Reference markers labelled for size in base pairs. **D**: Sanger sequence trace of a PCR products in Panel A (indicated by the black triangle). Sanger sequencing was carried out using the Applied Biosystems Big Dye Terminator version 3.1 to confirm PCR products as previously described. _17_ The Geneious^®^: Multiple Sequence Aligner tool was used to match the Sanger sequence of the sample with the predicted isoform as a reference sequence. E: BLAT alignment tool showing sequence match to *BRCA1* using the UCSC Genome Browser.

### BRCA1 isoform discovery and annotation

The first MinION sequencing run with R9 flow-cell chemistry using pooled RT-PCR products produced a total of 105,482 reads, of which 23,978 passed the quality control filters for 2D reads by Metrichor analysis. Of the total, 4,302 aligned to *BRCA1* in BLAT analysis, (and 54,505 aligned to another targeted gene sequence as part of a separate study), leaving 46,675 unaligned. The second MinION sequencing run produced a total of 12,022 reads, of which 995 passed the quality control filters for 2D reads by Metrichor analysis and 11,027 failed. As expected, the number of reads failing the 2D filter equated to 40-50% of output read numbers, presumably due to shearing of DNA by pipetting or incomplete ligation of hairpin adaptors, thus resulting in shorter reads lacking a complementary strand signal.

A total of 32 *BRCA1* isoforms (including full-length; Figure 2) were resolved with at least one sequencing read using a conservative mapping approach (Table 1). Of these, 20 isoforms have not been previously described. Of the 32 isoforms amplified by long-range RT-PCR, 23 lacked all or part of the largest *BRCA1* exon (exon 11; 3426 bases) (Table 1). Ten of these 23 isoforms contained a Δ11q splicing event rather than the complete skipping of exon 11. This results suggest that long-range RT-PCR assays were selective for shorter amplicons corresponding to smaller (<4kb) isoforms and that the sequencing results from the MinION may not be quantitative. The remaining nine of the detected transcripts were greater than 5kb in length and included the full length and Δ9,10 isoforms (Table 1), which have previously been shown to be ‘predominant’ transcripts in blood and breast cells using semi-quantitative measures.^9^ It is therefore possible that amplicon selection during long-range RT-PCR cycles may also have been influenced by the relatively high proportion of transcript levels in a pool of *BRCA1* expressed isoforms.

**Figure 2.**
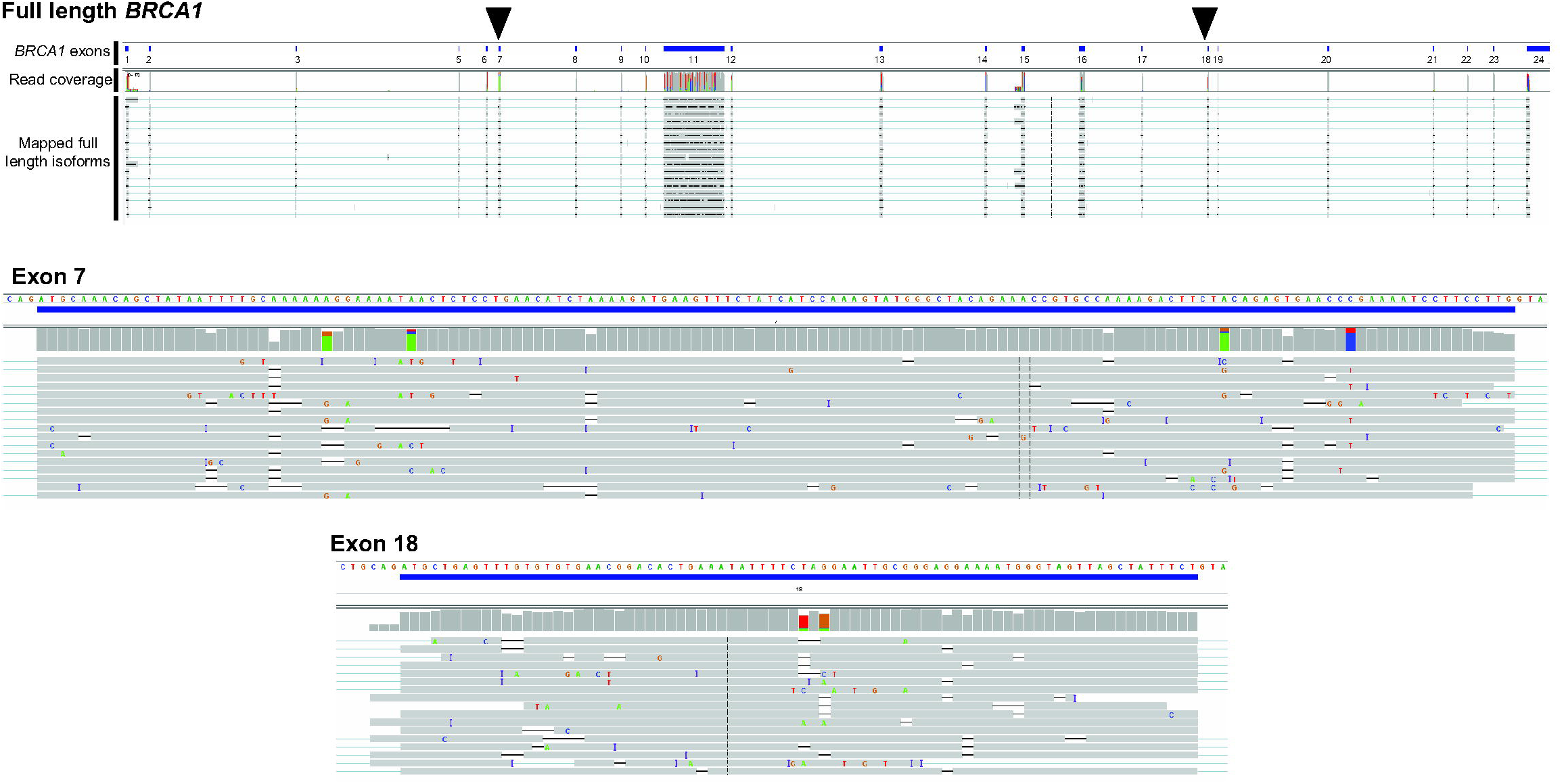
Sequencing of the full-length *BRCA1* transcript. Integrated Genome Viewer (IGV) screenshots are shown for the whole gene, along with close-up views of exons 7 and 18 (also highlighted by black triangles on full-length *BRCA1*). *BRCA1* exons are indicated and represented as blue solid rectangles. Each MinION sequence read with perfect homology to the reference sequence is shown in grey. Mismatches are shown in colour and indicated by base. The Oxford Nanopore MinION Genomic DNA Sequencing Kit (R9 flow cell chemistry) was used to prepare the DNA libraries according to the manufacturer’s instructions. Briefly, PCR products were purified then quantified using the Qubit^®^ Fluorometer (ThermoFisher Scientific) followed by end repair and dA tailing using the NEBNext Ultratm End Repair/dA-Tailing Module (New England BioLabs Inc). The DNA library entailing adaptor ligation and purification of double stranded DNA with hairpin adaptor was prepared using Nanopore Sequencing Kit SQK-NSK007 (R9 Version). The MinKNOW program was used for running the MinION for 48 hours. Additional sample mix was applied to the flow cell when the number of pores being used was less than 20, until the entire sample was used. The raw electrical signal was uploaded to Metrichor (version 1.107), using the 2D Basecalling RNN for SQK-NSK007 which returned basecall data in MinION fast5 file format. Sequence reads were mapped by Genomic Mapping and Alignment Program (GMAP)_18_ is using the linux command lines gmap -g [ReferenceSequence].fasta -f 2 -n 0 -t 16 [SequencesToAlign].fasta > [alignmentFile].gff3 as gmap -g BRCARD1 geneseq.fasta -f 2 -n 0 -t 16 all.fasta > all.gff3. These export output files as gff3.

**Table 1.** *BRCA1* isoforms identified by long-range PCR and nanopore sequencing

| Isoform designation | RNA | Size of isoform (bp) | Number of MinION reads | In-frame / Out-of-frame | Deleted protein domain(s)<sup>a</sup> | Previously described by Colombo *et al*<sup>b</sup> |
| --- | --- | --- | --- | --- | --- | --- |
| Δ11q | r.788_4096del | 3898 | 31 | In-frame | - | Yes |
| Δ9,10,11q | r.548_670del + r.788_4096del | 3775 | 27 | In-frame | - | Yes |
| Full length | - | 7207 | 17 | In-frame | - | Yes |
| Δ10-17 | r.594_5074del | 2726 | 14 | Out-of-frame | BCRT | No |
| Δ9,11q | r.548_593del + r.788_4096del | 3852 | 10 | Out-of-frame | - | Yes |
| 11Δ3110 | r.788_3897del | 4093 | 8 | In-frame | - | Yes |
| Δ1Aq,9,10,11q,12-15,16p | r.-25_-20del + r.548_670del + r.788_4804 | 3061 | 7 | In-frame | - | No |
| Δ14 | r.4358_4484del | 7080 | 6 | Out-of-frame | - | Yes |
| Δ10,11,17 | r.594_4096del + r.4987_5074del | 3616 | 5 | In-frame | BCRT | **No** |
| Δ1Aq,11Δ3110,Δ14p,15-17 | r.-25_-20del + r.788_3897del + r.4358_4360del + r.4485_5074del | 3498 | 5 | Out-of-frame | BCRT | **No** |
| Δ9,10 | r.548_670del | 7084 | 5 | In-frame | - | Yes |
| Δ1Aq,5q,11q,14,21 | r.-25_-20del + r.191_212del + r.788_4096del + r.4358_4484del + r.5278_5332del | 3688 | 5 | Out-of-frame | RING | **No** |
| Δ3 | r.81_134del | 7153 | 5 | Out-of-frame | RING | Yes |
| Δ9,10,14 | r.548_670del + r.4358_4484del | 6957 | 3 | Out-of-frame | - | **No** |
| Δ3-23 | r.81_5467del | 1820 | 3 | Out-of-frame | RING, BCRT | No |
| Δ2,9,10,11q,▼21 | r.-19_80del + r.548_670del + r.788_4096del + r.74421_74422ins74421+873_74421+1001 | 3676 | 3 | Out-of-frame (loss of coding start site) | RING | No |
| Δ9-11 | r.548_4096del | 3658 | 3 | In-frame | - | Yes |
| Δ2,9,10,11Δ3110,Δ14,20,22 | r.-19_80del + r.548_670del + r.788_3897del + r.4358_4484del + r.5194_5277del + r.5333_5406del | 3586 | 3 | Out-of-frame (loss of coding start site) | RING, BCRT | **No** |
| Δ11q,21 | r.788_4096del + r.5278_5332del | 3843 | 3 | Out-of-frame | BCRT | **No** |
| Δ11 | r.671_4096del | 3781 | 2 | In-frame | - | Yes |
| Δ3,11Δ3110 | r.81_134del + r.788_3897del | 4039 | 1 | Out-of-frame | RING | **No** |
| Δ9,10,21 | r.548_670del + r.5278_5332del | 7029 | 1 | Out-of-frame | BCRT | **No** |
| Δ21 | r.5278_5332del | 7152 | 1 | Out-of-frame | BCRT | Yes |
| Δ3,9,10,11q | r.81_134del + r.548_670del + r.788_4096del | 3721 | 1 | Out-of-frame | RING | **No** |
| Δ15-17 | r.4485_5074del | 6617 | 1 | Out-of-frame | BCRT | Yes |
| 11Δ3110,14,20-22 | r.788_3897del + r.4358_4484del + r.5194_5406del | 3753 | 1 | Out-of-frame | BCRT | No |
| Δ8,11 | r.442_547 + r.671_4096del | 3675 | 1 | Out-of-frame | - | No |
| 11Δ3110,Δ14 | r.788_3897del + r.4358_4484del | 3966 | 1 | Out-of-frame | - | **No** |
| Δ11q,19,20 | r.788_4096del + r.5153_5193_+ r.5194_5277del | 3773 | 1 | Out-of-frame | BCRT | No |
| Δ5,9,10,11q | r.135_212del + r.548_670del + r.788_4096del | 3697 | 1 | In-frame | RING | **No** |
| Δ3,9 | r.81_134del + r.548_593del | 7107 | 1 | Out-of-frame | RING | **No** |
| 11Δ3110,Δ21,23 | r.788_3897del + r.5278_5332del + r.5407_5467del | 3977 | 1 | Out-of-frame | BCRT | **No** |

Abbreviations: bp, base pairs; BCRT, BRCA1 C-terminal domain; RING, RING finger domain.
<sup>a</sup> Only clinically important BRCA1 protein domains are shown (ENIGMA BRCA1/2 Gene Variant Classification Criteria, Version 2.5 29th June 2017 https://enigmaconsortium.org/library/general-documents/). Position of amino acid residues denoting BRCA1 protein domains are as follows: RING 1-101 and BCRT, 1651-1863.<sup>b</sup> Novel isoforms that are created entirely from previously identified partial or whole exon skipping events are shown in bold.

Colombo et al previously characterized a total of 63 *BRCA1* alternative splicing events^9^, 17 of which were detected and further validated in this study. Eighteen of the 20 full-length novel isoforms identified by MinION sequencing were found to contain co-occurring exon skipping events (Table 1). Many of these events were found to co-occur, such as Δ(9,10), Δ11q, Δ(9,10,11q), and Δ3, have been previously identified in isolation using partial transcript analyses by Colombo *et al.^9^* Of the 20 novel isoforms discovered in this study, 14 were formed entirely from a combination of previously identified alternative splice junctions (Table 1). While these data show that many non-contiguous exon skipping events for *BRCA1* mRNA occur concurrently, this analysis was non-quantitative and therefore was unable to establish the relative levels of different transcripts.

Two novel isoforms identified, Δ10-17 (Supplementary Figure S3) and Δ3-23, show skipping of multiple contiguous exons generating out-of-frame coding sequences. We are unaware of previous studies that have implemented a PCR-based assay design that has encompassed exons 10-17 or exons 3-23. It is therefore not surprising that a long-range PCR-based approach has detected for the first time such isoforms. It is unlikely that Δ10-17 and Δ3-23 give rise to functional proteins as they lack the BRCA1 C-terminal (BCRT) domain (Table 1). Furthermore, the out-of-frame coding sequences for these isoforms suggest that they would be susceptible to nonsense mediated decay (NMD).^13^

The novel Δ10-17 isoform was selected for validation by Sanger sequencing as the isoform was common within the PCR amplicon library, and had a single junction which was relatively straightforward to amplify. RT-PCR assays using oligonucleotide primers targeting the exon 9-18 junction, followed by Sanger sequencing, confirmed the presence of this novel isoform (Figure 3).

**Figure 3.**
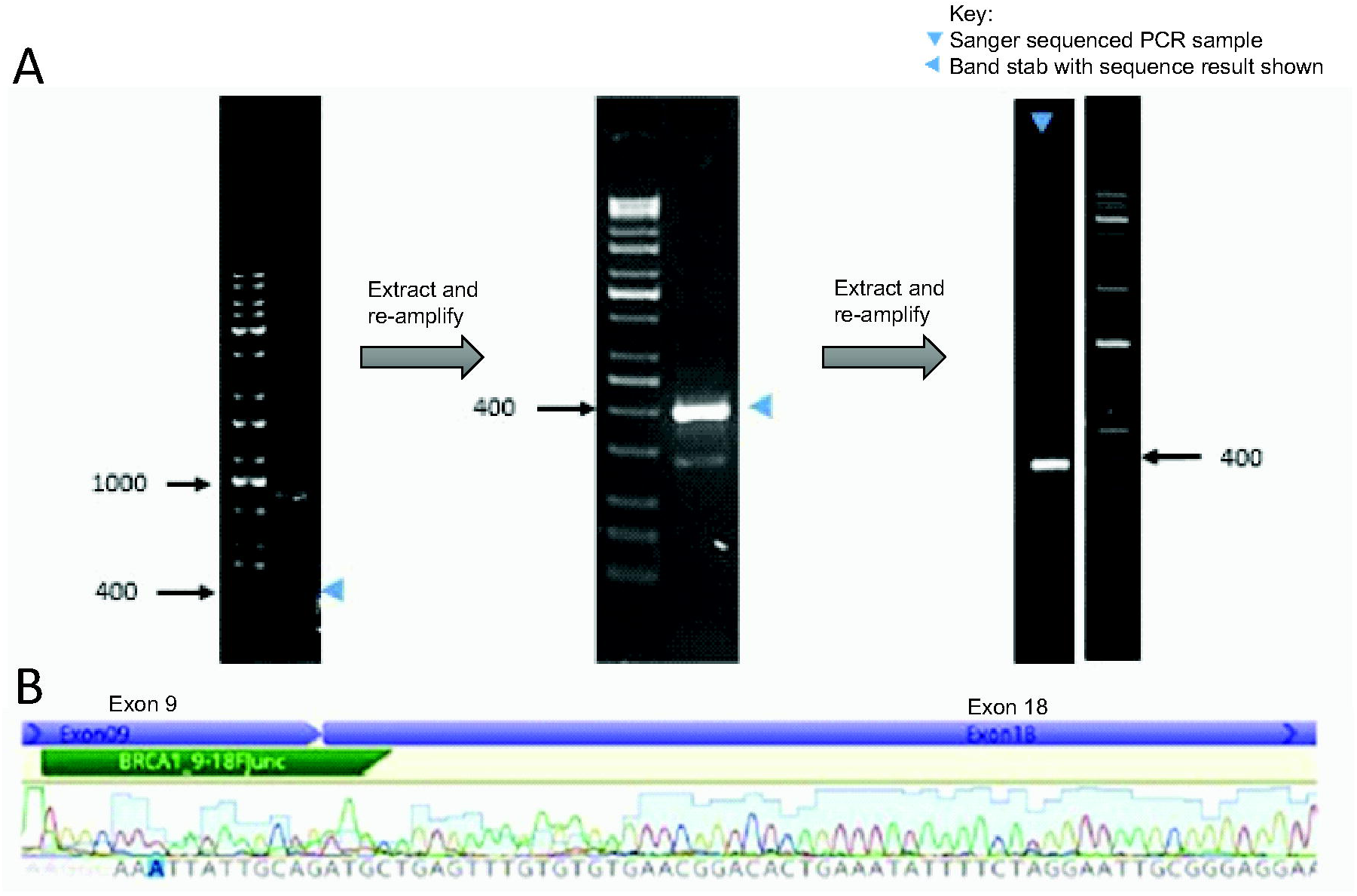
RT-PCR and Sanger sequencing confirmation of isoform Δ10-17. **A**. RT-PCR analysis of mRNA isolated from a cycloheximide treated LCL. To obtain a single isoform amplicon with sufficient DNA for sequence analysis, band extraction (shown by blue arrow heads) and re-amplification was carried out twice. The final 408bp product was analysed by Sanger sequencing. PCR reactions contained 1.3M betaine (Sigma-Aldrich), 1x KAPA long-range buffer, 1.75mM MgCl_2_, 0.5μM of each primer (BRCA1_9-18FJunc 5’-AGGCAAATTATTGCAGATGC-3’ and BRCA1_23RJunc 5’-ATGGAAGCCATTGTCCTCTG-3’), 300μM of KAPA, 200μM dNTP mix, and 0.5 units of KAPA Long Range HotStart. Thermal cycling conditions were 94°C for 4 minutes, followed by 35 cycles of 94°C for 30 seconds, primer annealing at one of a range of temperatures (56.4oC to 62.5oC; Supplementary Figure 2) for 30 seconds, and 68°C for 12 minutes, with a final extension of 72°C for 12 minutes. Primer annealing temperature is indicated above each lane. Reference markers labelled for size in base pairs. **B**. Sanger sequence trace of the exon 9-18 novel splice junction from the PCR product indicated in Panel A. Location of BRCA19-18 junction specific PCR primer (BRCA1_9-18FJunc) is indicated in green.

Together, these results suggest complexity in transcript structure that has not previously been described for *BRCA1.* Due to the potential error rate of the MinION (>10%)^14^, a higher read depth would increase the confidence in 19 of the 32 characterised transcripts represented by a relatively small number (n≤3) of reads, and this would be particularly important for potential splice shift events not previously identified.

### Co-occurring splicing events and interpretation for variant classification

Determining whether *BRCA1* transcripts lead to abnormal and potentially deleterious proteins requires knowledge about the structure of the coding isoforms. Sixteen of the 20 novel isoforms lacked sequences coding for the RING and/or BCRT domains, which have been previously shown to harbour amino acid residues of clinical importance^15^, although 14 of these 16 isoforms are out-of-frame and would therefore be susceptible to NMD. The remaining two isoforms (Δ(10,11,17) and Δ(5,9,10,11q)) are predicted to be in-frame and may potentially give rise to proteins lacking the clinically significant BCRT and RING domains, respectively.

We detected the predominant splicing events, Δ11q and Δ9,10, which individually would be predicted to generate modified in-frame transcripts that are not considered deleterious based on protein coding. However, our study has shown these splicing events can co-occur with skipping and insertion events causing out-of-frame coding in these isoforms (Table 1). Two examples were the Δ(9,10,21) and Δ(11q,21) isoforms which cause out-of-frame coding due to the exon 21 skipping event. Furthermore the combination of multiple out-of-frame exon skipping events can result in segments of the resulting transcript being in-frame. For example, Δ10,11 in the Δ(10,11,17) isoform is an out-of-frame deletion, but together with the out-of-frame Δ17 event, the isoform returns to being in-frame coding from exon 18 to exon 24. This transcript lacks sequence for the BRCT domain, suggesting that the isoform may avoid NMD and generate a protein that does not retain full BRCA1 function. Obtaining whole transcript information may therefore have important implications for interpreting the biological and clinical significance of spliceogenic variants.

While our work highlights that many *BRCA1* mRNA splicing events occur concurrently, our non-quantitative study was unable to establish the likelihood of such events being expressed in the same transcript. If future studies show that many of the detected *BRCA1* exon skipping events exclusively co-occur then the total number of isoforms might be lower than the number of events previously reported by Colombo et al.^9^ Such a finding would suggest similar splicing patterns may also exist for genes other than *BRCA1.* Further studies will therefore be required to measure the cellular levels of sequenced isoforms, and to investigate the possibility of further transcript complexity due to splice site shifts involving a small number of nucleotides. Such studies will require improved data analysis tools to take full advantage of the sequencing information generated, although we note that this field continues to advance as evidenced by a recent report by Hu et al^16^.

The ENIGMA Splicing Working Group previously led a multi-centre study which highlighted methodological issues that confounded the interpretation of splicing results^6^. A major reason for these issues was determined to be PCR assay design and the restrictive positioning of primers which prevented detection of additional naturally occurring isoforms. Our follow-up study has been successful in demonstrating the capability of the MinION device to characterize the exon structure of whole *BRCA1* transcripts. Together, our results highlight the potential of this technology to overcome limitations of traditional PCR-based techniques.

## Conclusion

Our study highlights complexity in *BRCA1* transcript structure that has not been described by previously reported studies. Assessment of whole *BRCA1* transcripts is now possible and has key implications for predicting translation frame of splicing transcripts, which is important for interpreting the clinical significance of spliceogenic variants. Future research is warranted to quantitatively assess full length *BRCA1* transcript levels, to detect additional novel isoforms involving small nucleotide shifts and to assess the application of nanopore sequencing for routine evaluation of potential spliceogenic variants. Furthermore, the application of the MinION or similar platforms may be extended to other disease associated genes to establish if they display similar complex splicing patterns to *BRCA1.*

## Acknowledgements

We thank the Cancer Society of New Zealand Canterbury/West Coast Division, and the Jim and Mary Carney Charitable Trust, for funding. LCW was supported by the Royal Society of New Zealand Rutherford Discovery Fellowship. ABS is supported by an Australian NMHRC Senior Research Fellowship. We wish to thank Heather Thorne, Eveline Niedermayr, all the kConFab research nurses and staff, the heads and staff of the Family Cancer Clinics, and the Clinical Follow Up Study (which has received funding from the NHMRC, the National Breast Cancer Foundation, Cancer Australia and the National Institute of Health (USA)) for their contributions to this resource, and the many families who contribute to kConFab. kConFab is supported by a grant from the National Breast Cancer Foundation, and previously by the National Health and Medical Research Council (NHMRC), the Queensland Cancer Fund, the Cancer Councils of New South Wales, Victoria, Tasmania and South Australia, and the Cancer Foundation of Western Australia.

